# Determinants of Murine Skin Microbiota Composition in Homeostasis and Wound Healing

**DOI:** 10.1101/2021.06.20.449197

**Authors:** Jack Galbraith, Julien M. D. Legrand, Nicholas Muller, Betoul Baz, Katie Togher, Nicholas Matigian, Seungha Kang, Sylvia Young, Sally Mortlock, Edwige Roy, Grant Morahan, Graeme Walker, Mark Morrison, Kiarash Khosrotehrani

## Abstract

Animal microbiota have complex interactions with hosts and environment that determines its composition. Yet the ability of hosts to determine their microbiota composition is less well studied. In this study, to investigate the role host genetics in determining skin microbiota, we used 30 different mouse strains from the recombinant inbred panel, the Collaborative Cross. Murine skin microbiota composition was strongly dependent on murine strain with > 50% of the variation explained by murine strain. In particular, a quantitative trait locus on chromosome 4 associates both with *Staphylococcus* abundance and principal-component multi-trait analyses. Additionally, excisional wound associated changes in microbiota composition were not uniform across mouse strains and were host-specific, the genetic background accounting for about 40% of the variation in microbiota. Genetic background also had the highest effect on the healing speed of wounds accounting for over 50% of the variation while mouse age and microbiota composition change accounted only for 20% and 5% of the healing speed despite reaching statistical significance. In conclusion, host genetics has a significant impact on the skin microbiota composition during both homeostasis and wound healing. These findings have long reaching implications in our understanding of associations between microbiota dysbiosis and disease.

## Introduction

In recent years, the microbiome has been implicated in a wide range of host responses relevant to homeostasis, and its dysbiosis can manifest in a wide range of immune- or metabolic-related diseases [1, 2]. The gut microbiome has been the most intensively studied and has advanced the concept of “community-based” interactions, associated with a range of conditions such as atopic dermatitis, obesity or mood disorders [3–5]. Although less studied, variations in the skin microbiome have been associated with episodes of atopic dermatitis [6], and the healing rate of leg ulcers, among other conditions [7, 8]. The skin microbiome has been shown to vary with anatomical location, patient age [9], and environmental cues such as humidity, temperature [10] or diet [2]. Overall, many environmental factors seem to determine the composition of the skin microbiota.

However, there are relatively few studies that provide a systematic assessment of how host genotype affects microbiome composition. Broader genome-wide studies utilising inter-cross mouse panel (BXD) have identified many host-specific quantitative trait loci (QTL) affecting gut microbiome composition, with candidate genes involving cytokines and toll-like receptor signalling [11]. In addition, investigation of the gut microbiota of mice from another advanced inter-cross model found 18 separate QTL associated with the relative abundance of one or more gut bacterial taxonomies [12]. Regarding skin, most studies to date have been limited to candidate-based approaches, such as how specific mutations affecting the serine protease matriptase can lead to a shift in the skin microbiota composition [13], or have described the skin microbial populations more generally [14–16]. In addition, many studies investigating skin wound microbiome have focused on non-healing chronic wounds such as diabetic foot ulcers and venous leg ulcers as opposed to the changes occurring in normal wound healing [17–19]. Whether host genetics also affects the wound microbiome, or alternatively, whether the environmental changes associated with wounding dominate over the effect of host genetic and more strongly affect microbial composition, are unknown.

Here we utilised a resource generated by a specialised breeding program, the Collaborative Cross (CC), which through a recombinant-inbreeding design, allows interrogation of complex traits and genetic pleiotropies [20, 21]. Using 30 different CC mouse strains, we investigated the host genetic contribution to mouse dorsal skin microbiota composition during homeostasis and wound healing. We found that the cutaneous microbiota composition dramatically differed between mouse strains, and that responses to wounding related to their microbiome composition were mouse strain-specific. Genome-wide association studies (GWAS) identified key QTL associated with specific bacterial taxa from normal skin. Overall our findings show that host genetics not only accounts for most of the observed variation in microbiome composition of normal skin, but also affects wound healing speed, while microbiota composition was found to only have a limited role in the latter process.

## Results

### Murine strain is a strong determinant of microbiota diversity

We collected swabs of mouse dorsal skin for 16S rRNA gene amplicon sequencing to investigate differences in microbiome composition between mouse strains. For each mouse, swabs were collected in a predefined area of the dorsal skin in a reproducible way. A negative swab exposed only to air was taken at the same time and sequenced with the OTU counts removed from downstream analysis as a normalising step.

We first compared the Shannon’s diversity metric as a measure of alpha-diversity (within sample) across 114 mice from 30 different mouse strains. The microbiota profiles from most mice fell within a relatively narrow range of Shannon’s diversity values (IQR=0.59, median=2.11), but communities in some mice clearly displayed less diversity and/or evenness (**Fig. 1a**). Although the intra-strain variation in Shannon’s diversity values was generally small, the inter-strain variation, as measured by Kruskal-Wallis testing was large and significant (*p*<0.0001). This analysis revealed that the large heterogeneity in overall mouse skin microbiota composition was strongly strain-dependent, which was further confirmed by PERMANOVA (R-squared > 0.5, *p*=0.001). We next asked whether specific microbial genera were associated specifically with lower or higher diversity compositions (‘ALDEx2’ R package). The abundances of both *Staphylococcus* and *Aerococcus* spp. significantly differed across the highest and lowest quartiles of Shannon’s diversity values, respectively (*Staphylococcus p*=0.0019, *Aerococcus p*=0.0028). A linear regression model using Shannon’s diversity and the centred-log ratio (CLR) transformation of the abundance values for *Staphylococcus* and *Aerococcus* across all mice showed there was an inverse relationship of both genera with the Shannon’s diversity value of mouse skin microbiome (**Supp. 2b**). This highlighted the importance of the abundance of these two genera on the overall skin microbiota composition.

**Figure 1.**
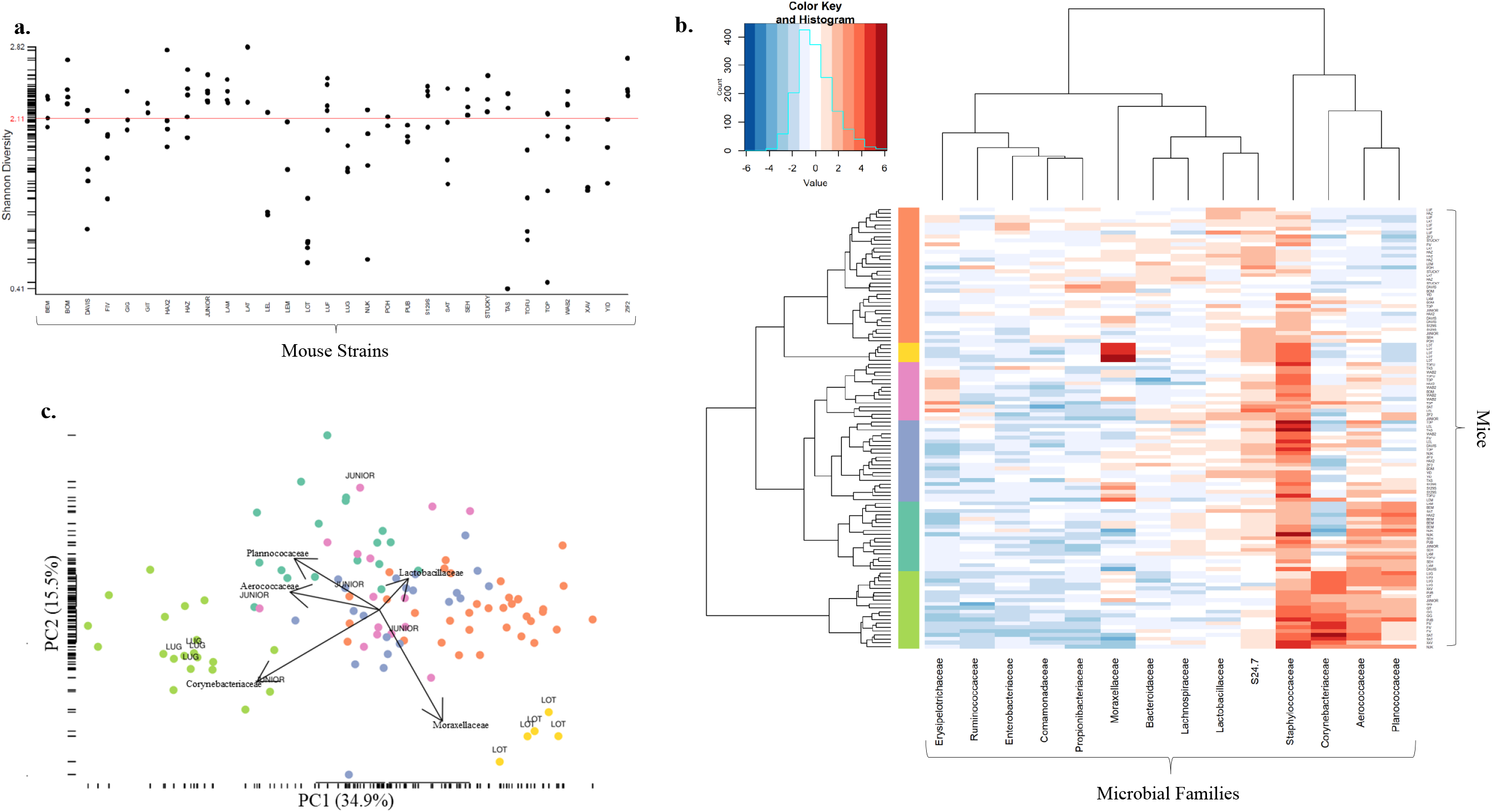
An overview of the cutaneous microbiota across 30 different strains of mice from the Collaborative Cross. (**a**) Scatterplot of alpha diversities for each sample, grouped by mouse strain. Most mice show similar alpha-diversities and low intra-strain variability. Diversity of the skin microbiome shows strong strain dependent differences (Kruskal-wallis, *p*-value<0.0001). (**b**) Hierarchical clustering of microbiota composition (centred-log ratio read counts) of healthy dorsal skin swabs. Many mice cluster strongly within their respective strain (**c**) Principal component analysis of centred-log ratio read counts further highlights the differences in abundance of microbiota families between mice.

Next, we determined a core skin microbiome and used different levels of bacterial taxonomies to identify their representation in at least 50% of samples at 0.1% or greater relative abundance. Importantly, unbiased agglomerative hierarchical clustering of animals based on their skin microbial composition (family level) [22, 23] demonstrated that mice clustered within their respective strains (**Fig. 1b**). Similarly, principle component analysis of microbiota composition revealed the grouping of individual mice based on clusters (**Fig. 1c**). While *Staphylococcaceae* was the most prevalent bacterial family in the strains of mice examined, and thereby a member of the core microbiome, there was also a group of mouse strains that clustered based on their possession of a relatively high abundance of *Corynebacteriaceae* (**Fig. 1b & Fig. 1c**). Although mouse age significantly affected skin microbiome profile, it was found to explain only a minimal amount of the variance observed across mice (R-squared < 0.02). Taken together, these findings strongly confirmed the effect of murine strain (host genotype) as a key determinant of the dorsal skin microbiome.

### Skin microbiota composition associates with murine strain specific genomic loci

We next examined whether specific loci in the mouse genome could be associated with the dorsal skin microbiome composition. The centred log-ratio transformation of the abundance values for *Staphylococcus* was used as a “trait” for host genome-wide association analysis and identified a genome-wide significant region on mouse chromosome (Chr) 4 between 129.75-130.95 megabase pairs (Mbps) (LOD Score: 10, **Supp. 1a**). Several different loci, albeit with weaker (suggestive) LOD scores were identified using values for *Aerococcus* (Chr 13, 108.70-113.51 Mbps, LOD Score: 6, **Supp. 1b**), Shannon’s diversity scores (Chr 15, 3.20-7.40 Mbps, LOD Score: 8, **Supp. 1c**) or principal component analysis (PCA)-based GWAS [24] (**Supp. 1d-e**). Interestingly, the same peak on Chr 4 identified for *Staphylococcus* was also recovered using the PCA-based GWAS (**Supp. 1e**). A detailed analysis of the founder haplotypes for the Chr 4 candidate region identified the WSB and PWK founder alleles associated with a relatively low abundance of *Staphylococcus* in skin microbiome, and the CAST founder allele with highest relative abundances. We next examined this region of interest for any genes harbouring specific polymorphisms in the founder haplotypes above, by using the Sanger Mouse Genome Project SNP query website. Here, we identified *Ptafr* as a candidate gene of interest, on the basis that its product may affect the immune response to pathogens [25, 26], but is also known to act directly on the wound healing process [27], and can affect skin inflammation [28]. An alternative candidate is *Smpdl3b, Sphingomyelin Phosphodiesterase Acid-Like 3B*, which is associated with inflammation via negative regulation of toll-like receptor signalling *in vivo* [29]. A full list of genes containing haplotype specific SNP is provided in **Suppl. Table.1**. Overall, this strong effect of murine strain, as well as the association with plausible candidate genes, strongly supports the importance of host genetics in determining the skin microbiome during homeostasis.

### The microbiomes of early stage wounded and normal skin retain host-strain specificity

We next investigated whether wounding, considered a major environmental stress, would elicit stronger effects on the microbiome than host genotype. If skin injury and wound healing had strong effects on the microbiome, one would expect the wound microbiome to be convergent across the different mouse strains. Alternatively, given the exposure to faecal material in the cage bedding, one would expect the wound and faecal microbiome to converge. To that end, we performed PCA and hierarchical clustering on the combined datasets from unwounded skin, day (D) 3 wounds, and the faecal microbiome samples of 70 mice representing the different haplotypes. We found that the wound microbiome at D3 retained its similarity to the community present on unwounded skin, and that the faecal microbiomes were clearly separable from the skin and wound microbiomes using PCA and hierarchical clustering **(Fig. 2a-b)**. The bacterial families *Propionibacteriaceae* and *Staphylococcaceae* were discriminatory for both the unwounded skin and wound microbiomes at D3, compared to faecal samples, and *Bacteroidales* family S24-7 were in much greater abundance in faecal samples (**Supp. 2a**). These findings clearly showed that wound microbiota composition does not assimilate with the faecal microbiota despite the large environmental exposure to faeces.

**Figure 2.**
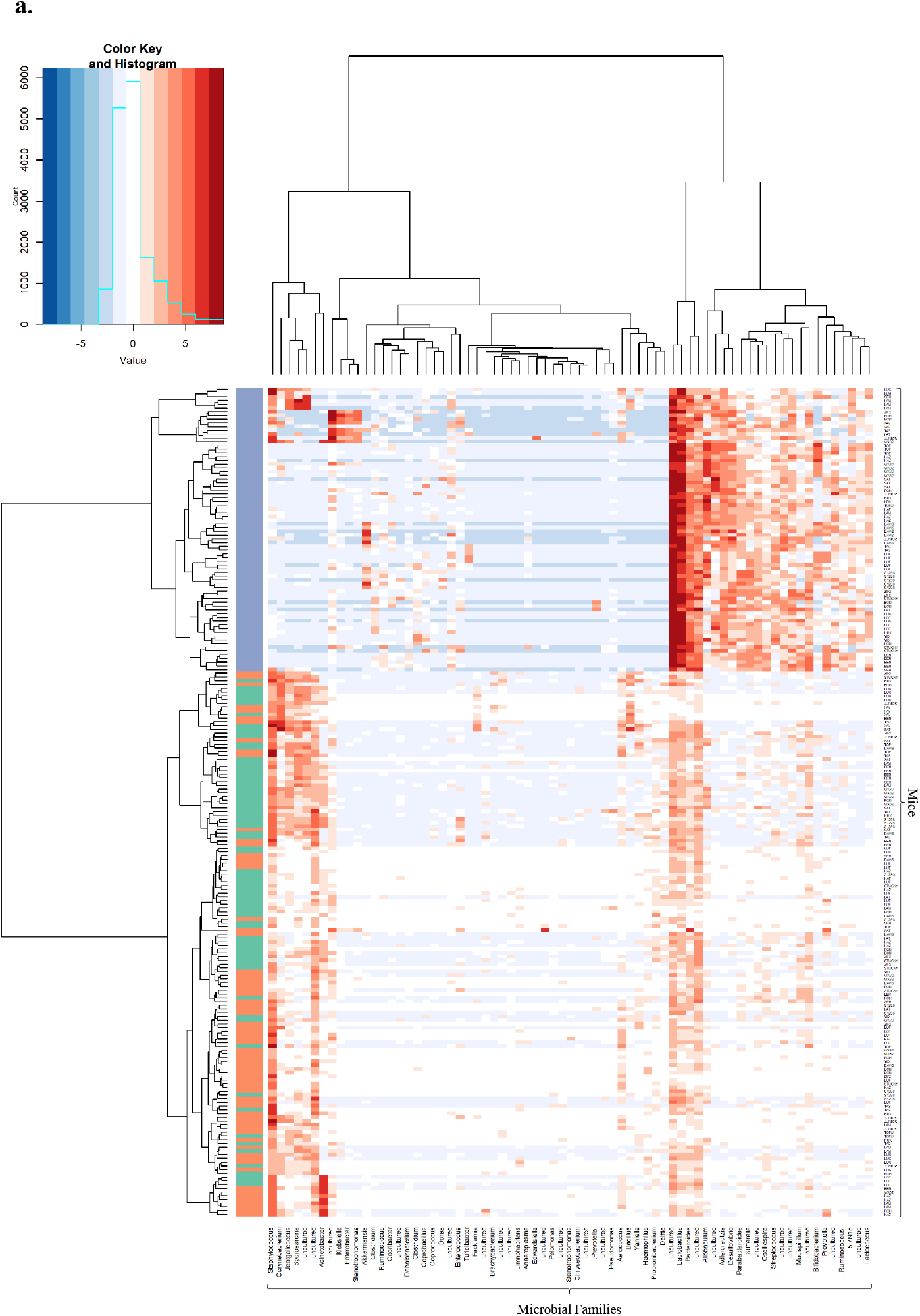
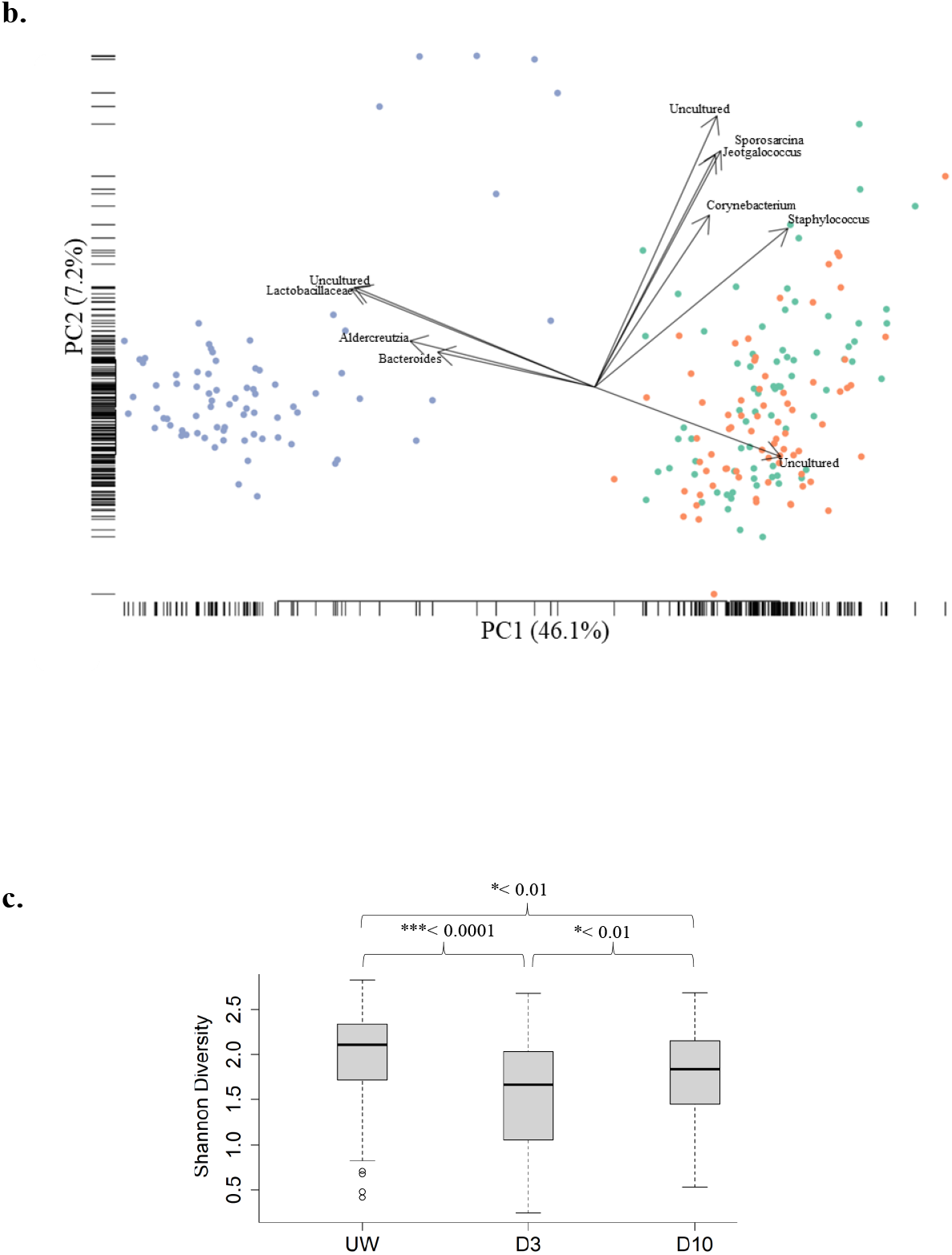
Comparison of diversity and composition of faecal, unwounded and wounded skin microbiota. (**a**) Hierarchical clustering heatmap of centred-log ratio read counts from all 3 sample microbiota types (faecal = blue, unwounded = green, day-3 post-wounding = orange). (**b**) Principal component analysis of combined sample sets (faecal = blue, unwounded = green, day-3 post-wounding = orange) at the genus level. Day-3 post-wound microbiota cluster with unwounded microbiota samples and separately from faecal samples (**c**) Skin microbial diversity at different time-points (Shannon alpha diversity index). Box plot displaying the overall alpha diversity from unwounded to day-3 and day-10.

While the D3 wound microbiomes retained features of normal skin, they displayed a significant decline in diversity (**Fig. 2c**, Mann-Whitney, *p*-value<0.0001, 0.788 median fold-change in Shannon’s diversity across all mice). These declines in microbial diversity of the wounds were only transient and recovered significantly by D10 post-wounding (Kruskal-Wallis testing *p*<0.01, 1.1 median fold-change in Shannon’s diversity across all mice), but remained significantly lower than the original unwounded skin at this time-point (*p*<0.01, 0.8 median fold-change unwounded to D10 Shannon’s diversity).

Given the broad reduction in diversity we next wondered whether there was a uniform change in microbiota after wounding. The hierarchical clustering (**Fig. 3a**) showed that although there was a general trend of increased *Staphylococcaceae* and *Corynebacteriaceae* relative abundances between D3-wounded and unwounded skin (FDR-adjusted, *p*<0.1, ALDEX2) in many mice, some strains, GIG and WAB2, showed a decrease in these families. This clearly highlighted that the changes of microbiota composition in response to wound healing were not universal and varied across murine strains.

**Figure 3.**
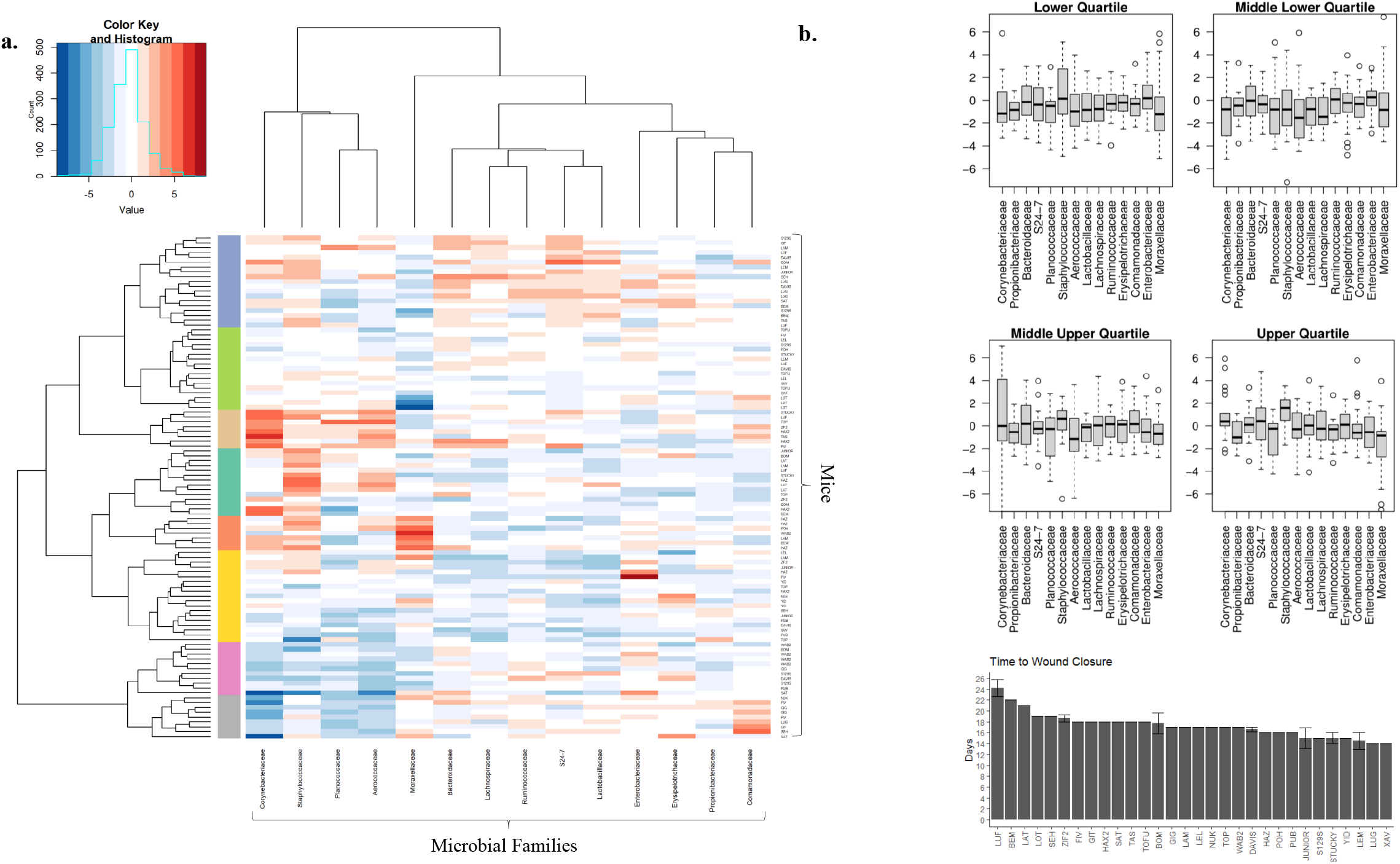
Microbiota composition changes from unwounded skin to day 3 post-wounding. (**a**) Hierarchical clustering of microbiota abundance change from unwounded to D3 wounds. Taking the difference between day 3 post-wounding and unwounded centred-log ratio matrices shows the relative increase/decrease in microbial abundance during the early stages of wound healing. (**b**) Boxplots of bacterial family centred-log ratio abundance changes during wound healing across all mice, grouped by quartiles of mouse healing speed (Top and Middle). Days to full wound closure across all mouse strains (Bottom).

To investigate the relative contributions to changes in microbiota composition during wound healing, a PERMANOVA analysis was performed with the variables being mouse strain, wound time-point (unwounded or D3) and mouse age (days). The strain of mouse had the largest effect on overall microbiome compositional changes during wound healing (~23% variance explained), followed by the interaction between mouse strain and wound time-points (~17% variance explained). While statistically significant, age and wound time-point alone accounted for only a small amount of variance (<8% combined). This once again strongly supports the effect of host genotype on both the unwounded and wounded skin microbiomes.

### Microbiome composition had minimal effect on wound healing rates

Given the variation between mice in their D3 post-wounding microbial composition and the microbiota differences from unwounded skin, we investigated whether these parameters were associated with wound healing. The time to wound closure for individual mice was determined and used to group these into quartiles representing fast to slow healers. There were large variations within each healing quartile, both positively and negatively, in the relative abundances of different bacterial families present in D3 wound microbiomes compared to unwounded skin (**Fig. 3b**). However, these changes in the skin/wound microbiomes during early wound healing, and the faecal microbiome composition of the host animal each accounted for ~5% of the variance in wound healing speeds when modelled separately by principal component regression. In contrast, PEMANOVA analysis showed mouse strain (genotype) accounted for over 50% of the variance in wound healing rate (*p*<0.001) while the age of mice explained 20% variance (*p*-value<0.001). Age is already recognised to be a significant factor in the regenerative ability of mice [30, 31]. Our results not only confirm the association between mouse age and wound healing speed, but the limited impact of age on skin microbiome composition suggests the impacts of age on wound healing speed are host-rather than microbiome-related.

## Discussion

The skin microbiome has been associated with the onset and/or progression of many skin disorders [5–7, 14, 17, 19, 32]. Many studies have revealed how environmental factors such as temperature, humidity, sun exposure, body site, or host factors such as age and diet can affect the skin microbiome [5, 9, 10, 33]. These collective findings have led many to infer that wounding - which is an extreme form of environmental insult - results in a stereotypical change in the composition of the microbiome at the wound site, principally via increases in the relative and total abundances of *Staphylococcus* and Gram-negative bacteria [32, 34]. In contrast, very little is known about whether and how host genotype affects the skin microbiome, and further, to what extent differences in wound healing speed are attributable to host-driven processes either directly, or indirectly, via the skin microbiome. Using the collaborative cross genetic resource, we conducted a large and systematic study to evaluate the importance of genetic background on murine skin microbiome composition and its impact on wound healing. Our findings highlight the strong association of host genetics with microbiome, even in the presence of a major environmental factor such as wounding. The impact of wounding and exposure to faeces did not induce uniform changes across all strains with host genetic remaining the dominant factor in microbiota variation. Finally, microbiota composition and change was shown to only minimally affect wound healing speed as opposed to genetic background.

Importantly, all the animals used in this study were born in the same animal facility; then all shipped and subsequently housed in a different facility and provided access to the same food and water sources. While it is not possible to completely rule out any housing effects across all 30 strains of mice, other studies have revealed that compared to host genotype, the contributions from caging and legacy effects on the variations in the gut microbiome are small [35]. Additional studies of skin bacterial populations in various mammals and amphibians further supports host taxonomy as a greater determining factor of the cutaneous microbiome than environment [36–38]. Mice of the same species kept in laboratory conditions or wild mice possess largely similar skin microbiome [37], whereas different species of amphibians co-habiting the same pond show divergent skin microbiomes [36]. Clearly across many species, the underlying genetics of the host have a significant determining effect on the overall skin microbiome. Our analysis was limited to the most abundant “core” bacterial families whereas others have suggested that differences in microbiomes between groups, such as diabetics and controls, is driven primarily through very low abundant bacteria [17]. For other aspects of our study, housing effects are largely unimportant such as or association analyses of microbiome with wound healing, as the regression analysis should identify associations with wound healing speeds regardless of whether these are due to genetic or environmental determinants. Similarly, many of the QTL identified in the GWAS are due to multiple mouse strains that share the same haplotype for that region of the genome but were not caged together.

In that context, the genome-wide associations identified key host loci predictive of specific skin bacterial taxa and/or microbiome composition. Indeed, specific bacterial genera (*Staphylococcus* and *Aerococcus*) and Shannon diversity scores of the skin microbiome from individual mice could be used as a “trait” in genome wide association and linkage studies. In particular, these analyses defined a ~1 Mbp region on Chr 4 to be strongly associated with these traits. Whilst this list provides only candidates that are associated with the “traits”, it does include intuitive genes such as those affecting host innate and adaptive immunity, for example *Ptafr*, or *Smpdl3b*. While the true role of these candidate genes needs experimental validation, it is plausible that SNP variations in any of these candidate genes may elicit differential immune or dermal niche alterations that affect the relative abundance of *Staphylococcus* and/or *Aerococcus* on skin.

In a clinical setting, it has been thought that skin wounds are characterised by major colonisation of Gram-negative bacteria, such as *Pseudomonas aeruginosa* [34], and often display an over-representation of Staphylococci [32]. Here, we report that wound microbiome remains highly variable in its composition across different murine strains and there was no homogenous change across all mice. Inclusion of stool samples allowed us to show that D3 post-wound skin retains a skin-like microbiota phenotype. This is unexpected given the extreme barrier function dysregulation as well as the relative contamination of cage bedding with faeces. Considering the site specificity and stability of the human microbiome [9], it is possible that the transcriptomic environment of skin wounds drives the microbiota to retain its skin-like features.

Our study highlights the difficulty of drawing conclusions about microbial associations with wound healing outcomes across studies that have used a single background mouse model of microbiota changes. Studies of chronic wounds in patients show relatively little difference in the more abundant genera between healing and non-healing wounds requiring the identification of specific species of bacteria to determine wound outcomes [39]. In line with these results, we were unable to identify any statistically significant, or suggestive, abundant microbial families associated with healing speed in mice. We did however find a decline in diversity as observed in eczema flares, and non-healing diabetic foot ulcers [14, 17].

In conclusion, we report that skin murine microbiota and its changes upon wounding are strongly determined by host genetics and the abundance of specific microbial families can be determined by precise loci in the murine genome. Moreover, the wound microbiome plays a minimal role in the healing rate and is mostly a reflection of the host genetic background. These findings have far reaching implications for the design of further studies on the role of the microbiome on health and disease as well as the use of probiotics in a clinical setting.

## Materials & Methods

### Mice

All mice from the collaborative cross (n=114) were housed in the UQ Centre for Clinical Research Animal Facility. All animal experimentation was conducted in accordance with institutional ethical requirements and approved by the University of Queensland Animal Ethics Committee. Only female mice were used with the number of mice per strain varying due to availability with a minimum of 2 for only 3 strains. Age of mice were recorded in **Supp.Table.2**.

### Collection

Mice were anaesthetized with 2% isoflurane and a sterile rayon swab moistened with TE buffer was used to collect a microbiota sample from a 1.5×1.5cm of dorsal skin, followed by full-thickness excisional wounds of the same area. At this initial timepoint a fresh stool sample was collected. Additionally, a negative control swab was exposed to air in the same room as the wounding procedures immediately prior to wounding. This sample was sequenced and a total of 5 OTU counts were retrieved. These were subsequently subtracted from all other samples as a normalising procedure. Additional swabs were taken at days 3 and 10 post-wounding. All samples were stored in 2ml of TE buffer at −80°C for later sequencing of the V6-V8 hypervariable region of the 16S rRNA gene using the universal microbial primers from Illumina.

Details of OTU annotation, and analyses as well as genome wide association studies are in the supplemental methods.

## Data availability statement

OTU tables are available in supplemental materials. All sequencing data is available on request until deposition is finalised.

## Acknowledgements

We would like to thank Geniad Pty Ltd for the generous provision of CC mice. This work was supported by funding from the National Health and Medical Research Council of Australia (APP1082438). KK is supported by a Queensland government, Advance Queensland Research Fellowship.

## Author contributions

Conceptualization: KK, MM, GW, GM; Data Curation: JL, BB, NM, ER, GM; Formal Analysis: JG, NM, KT, SK, SY, NM, SM; Funding Acquisition: GW, KK; Investigation: JL, BB, NM, ER, GM, GW, KK; Methodology: JG, SK, SM; Project Administration: KK, MM, GW; Resources: GM, MM, KK; Software: JG, NM, KT, SK, SM; Supervision: KK, MM, GW; Validation: JG, NM, KT, SK, SM; Visualization: JG, KK, SK.; Writing – JG, KK; Writing - Review and Editing: JG, KK, ER, MM, SK and all other authors

## References

1. Catinean, A., et al., Microbiota and Immune-Mediated Skin Diseases-An Overview. Microorganisms, 2019. 7(9).

2. Schoeler, M. and R. Caesar, Dietary lipids, gut microbiota and lipid metabolism. Rev Endocr Metab Disord, 2019. 20(4): p. 461–472.

3. Forsythe, P., et al., Mood and gut feelings. Brain Behav Immun, 2010. 24(1): p. 9–16.

4. Kobyliak, N., O. Virchenko, and T. Falalyeyeva, Pathophysiological role of host microbiota in the development of obesity. Nutr J, 2016. 15: p. 43.

5. Pessi, T., et al., Interleukin-10 generation in atopic children following oral Lactobacillus rhamnosus GG. Clin Exp Allergy, 2000. 30(12): p. 1804–8.

6. Kong, H.H., et al., Temporal shifts in the skin microbiome associated with disease flares and treatment in children with atopic dermatitis. Genome Res, 2012. 22(5): p. 850–9.

7. Min, K.R., et al., Association between baseline abundance of Peptoniphilus, a Gram-positive anaerobic coccus, and wound healing outcomes of DFUs. PLoS One, 2020. 15(1): p. e0227006.

8. Wolcott, R.D., et al., Evaluation of the bacterial diversity among and within individual venous leg ulcers using bacterial tag-encoded FLX and titanium amplicon pyrosequencing and metagenomic approaches. BMC Microbiol, 2009. 9: p. 226.

9. Grice, E.A., et al., Topographical and temporal diversity of the human skin microbiome. Science, 2009. 324(5931): p. 1190–2.

10. Grice, E.A. and J.A. Segre, The skin microbiome. Nat Rev Microbiol, 2011. 9(4): p. 244–53.

11. McKnite, A.M., et al., Murine gut microbiota is defined by host genetics and modulates variation of metabolic traits. PLoS One, 2012. 7(6): p. e39191.

12. Benson, A.K., et al., Individuality in gut microbiota composition is a complex polygenic trait shaped by multiple environmental and host genetic factors. Proc Natl Acad Sci U S A, 2010. 107(44): p. 18933–8.

13. Scharschmidt, T.C., et al., Matriptase-deficient mice exhibit ichthyotic skin with a selective shift in skin microbiota. J Invest Dermatol, 2009. 129(10): p. 2435–42.

14. Byrd, A.L., et al., Staphylococcus aureus and Staphylococcus epidermidis strain diversity underlying pediatric atopic dermatitis. Sci Transl Med, 2017. 9(397).

15. Grice, E.A., et al., A diversity profile of the human skin microbiota. Genome Res, 2008. 18(7): p. 1043–50.

16. Oh, J., et al., Temporal Stability of the Human Skin Microbiome. Cell, 2016. 165(4): p. 854–66.

17. Gardiner, M., et al., A longitudinal study of the diabetic skin and wound microbiome. PeerJ, 2017. 5: p. e3543.

18. Redel, H., et al., Quantitation and composition of cutaneous microbiota in diabetic and nondiabetic men. J Infect Dis, 2013. 207(7): p. 1105–14.

19. Loesche, M., et al., Temporal Stability in Chronic Wound Microbiota Is Associated With Poor Healing. J Invest Dermatol, 2017. 137(1): p. 237–244.

20. Churchill, G.A., et al., The Collaborative Cross, a community resource for the genetic analysis of complex traits. Nat Genet, 2004. 36(11): p. 1133–7.

21. Ferguson, B., et al., Melanoma susceptibility as a complex trait: genetic variation controls all stages of tumor progression. Oncogene, 2015. 34(22): p. 2879–86.

22. Murtagh, F., Legendre, P., Ward’s Hierarchical Agglomerative Clustering Method: Which Algorithms Implement Ward’s Criterion? Journal of Classification, 2014. 31: p. 274–295.

23. Ward, J.H., Hierarchical Grouping to Optimize an Objective Function. Journal of the American Statistical Association, 1963. 58: p. 263–244.

24. Zhang, W., et al., PCA-Based Multiple-Trait GWAS Analysis: A Powerful Model for Exploring Pleiotropy. Animals (Basel), 2018. 8(12).

25. Shukla, S.D., et al., Platelet activating factor receptor: gateway for bacterial chronic airway infection in chronic obstructive pulmonary disease and potential therapeutic target. Expert Rev Respir Med, 2015. 9(4): p. 473–85.

26. Iovino, F., et al., Signalling or binding: the role of the platelet-activating factor receptor in invasive pneumococcal disease. Cell Microbiol, 2013. 15(6): p. 870–81.

27. Stafforini, D.M., et al., Platelet-activating factor, a pleiotrophic mediator of physiological and pathological processes. Crit Rev Clin Lab Sci, 2003. 40(6): p. 643–72.

28. Sahu, R.P., et al., Topical application of a platelet activating factor receptor agonist suppresses phorbol ester-induced acute and chronic inflammation and has cancer chemopreventive activity in mouse skin. PLoS One, 2014. 9(11): p. e111608.

29. Heinz, L.X., et al., The Lipid-Modifying Enzyme SMPDL3B Negatively Regulates Innate Immunity. Cell Rep, 2015. 11(12): p. 1919–28.

30. Velarde, M.C., et al., Pleiotropic age-dependent effects of mitochondrial dysfunction on epidermal stem cells. Proc Natl Acad Sci U S A, 2015. 112(33): p. 10407–12.

31. Keyes, B.E., et al., Impaired Epidermal to Dendritic T Cell Signaling Slows Wound Repair in Aged Skin. Cell, 2016. 167(5): p. 1323–1338 e14.

32. Shettigar, K. and T.S. Murali, Virulence factors and clonal diversity of Staphylococcus aureus in colonization and wound infection with emphasis on diabetic foot infection. Eur J Clin Microbiol Infect Dis, 2020. 39(12): p. 2235–2246.

33. Kong, H.H., Skin microbiome: genomics-based insights into the diversity and role of skin microbes. Trends Mol Med, 2011. 17(6): p. 320–8.

34. Yung, D.B.Y., K.J. Sircombe, and D. Pletzer, Friends or enemies? The complicated relationship between Pseudomonas aeruginosa and Staphylococcus aureus. Mol Microbiol, 2021.

35. Khan, A.A., et al., Polymorphic Immune Mechanisms Regulate Commensal Repertoire. Cell Rep, 2019. 29(3): p. 541–550 e4.

36. McKenzie, V.J., et al., Co-habiting amphibian species harbor unique skin bacterial communities in wild populations. ISME J, 2012. 6(3): p. 588–96.

37. Belheouane, M., et al., Assessing similarities and disparities in the skin microbiota between wild and laboratory populations of house mice. ISME J, 2020. 14(10): p. 2367–2380.

38. Ross, A.A., et al., Comprehensive skin microbiome analysis reveals the uniqueness of human skin and evidence for phylosymbiosis within the class Mammalia. Proc Natl Acad Sci U S A, 2018. 115(25): p. E5786–E5795.

39. Verbanic, S., et al., Microbial predictors of healing and short-term effect of debridement on the microbiome of chronic wounds. NPJ Biofilms Microbiomes, 2020. 6(1): p. 21.

