## Supplemental_Figure_1 for "Determinants of Murine Skin Microbiota Composition in Homeostasis and Wound Healing"

### Supplemental 1.

a.

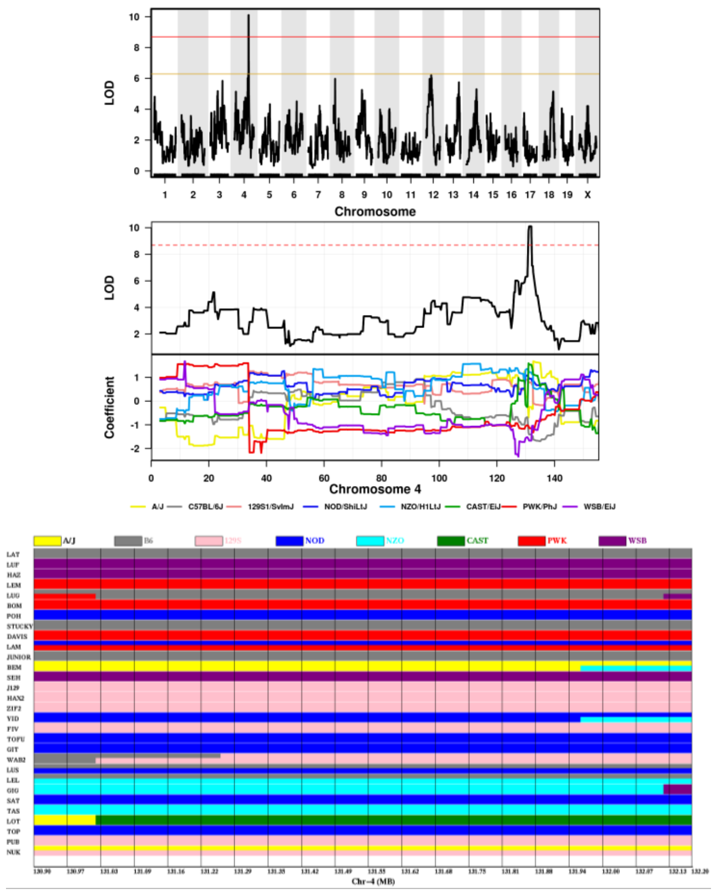


b.

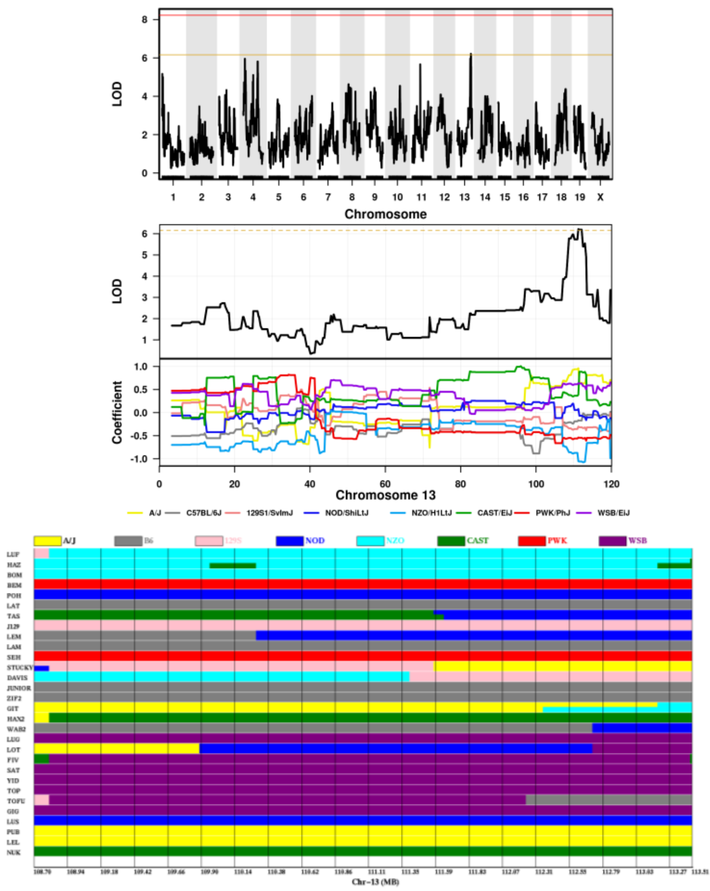


c.

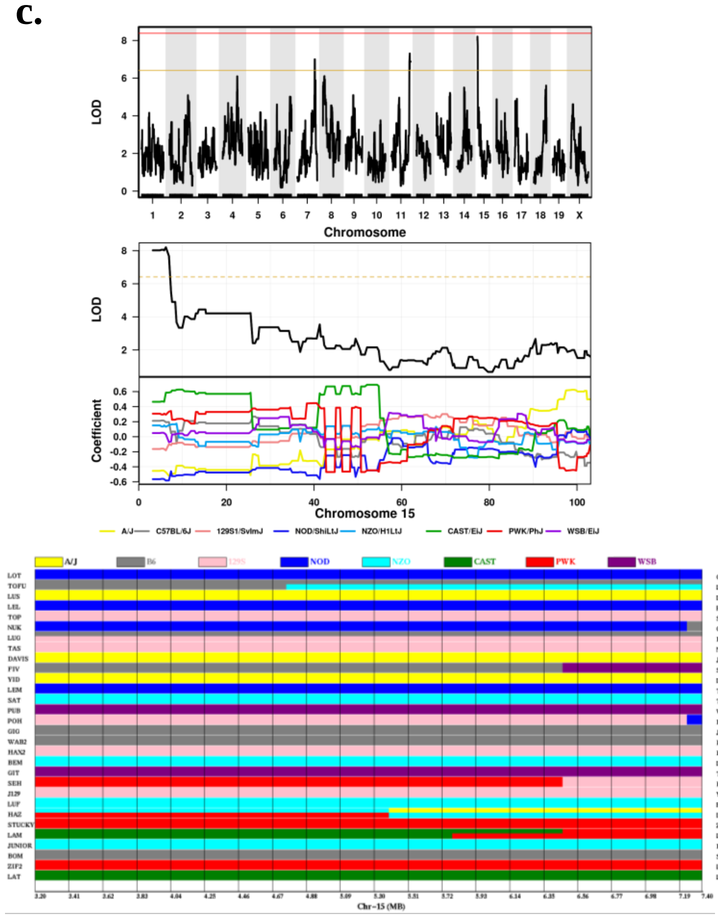


d.

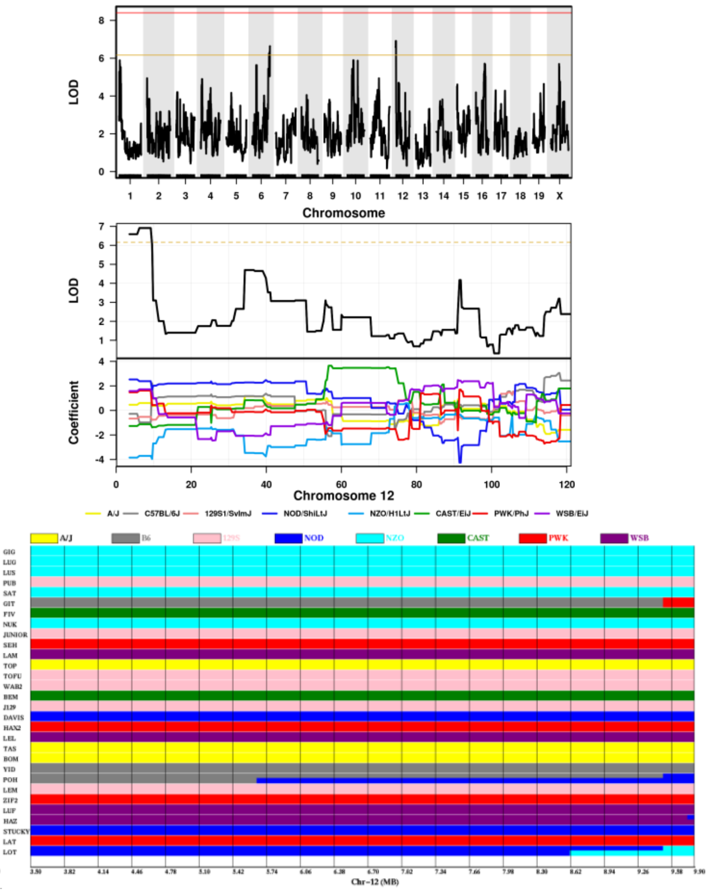

### Supplemental 1. Cont.

e.

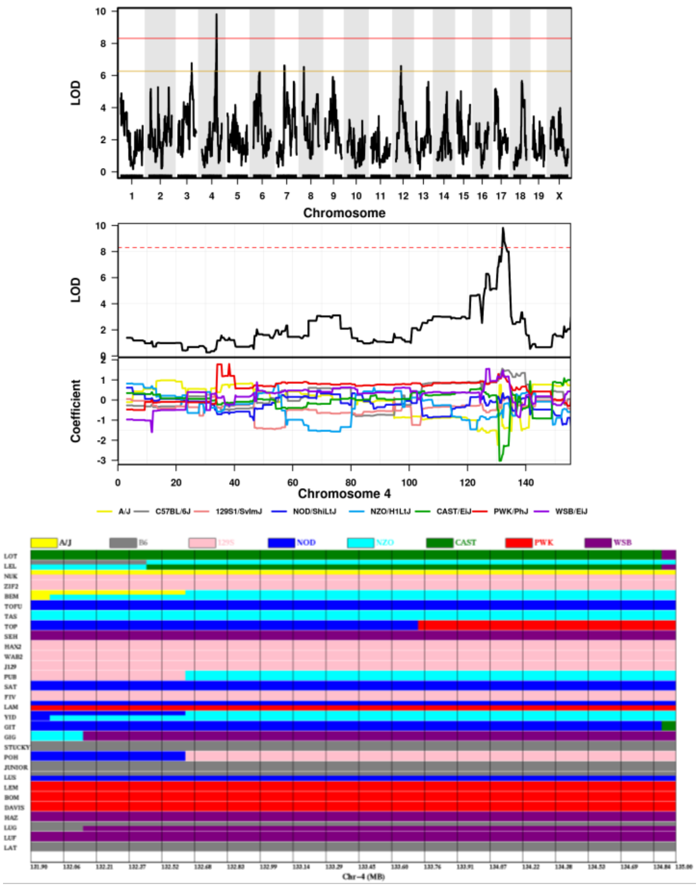

### Supplemental 2.

**a.** Log-proportional abundances of all bacterial families (including non-core) (70 mice)

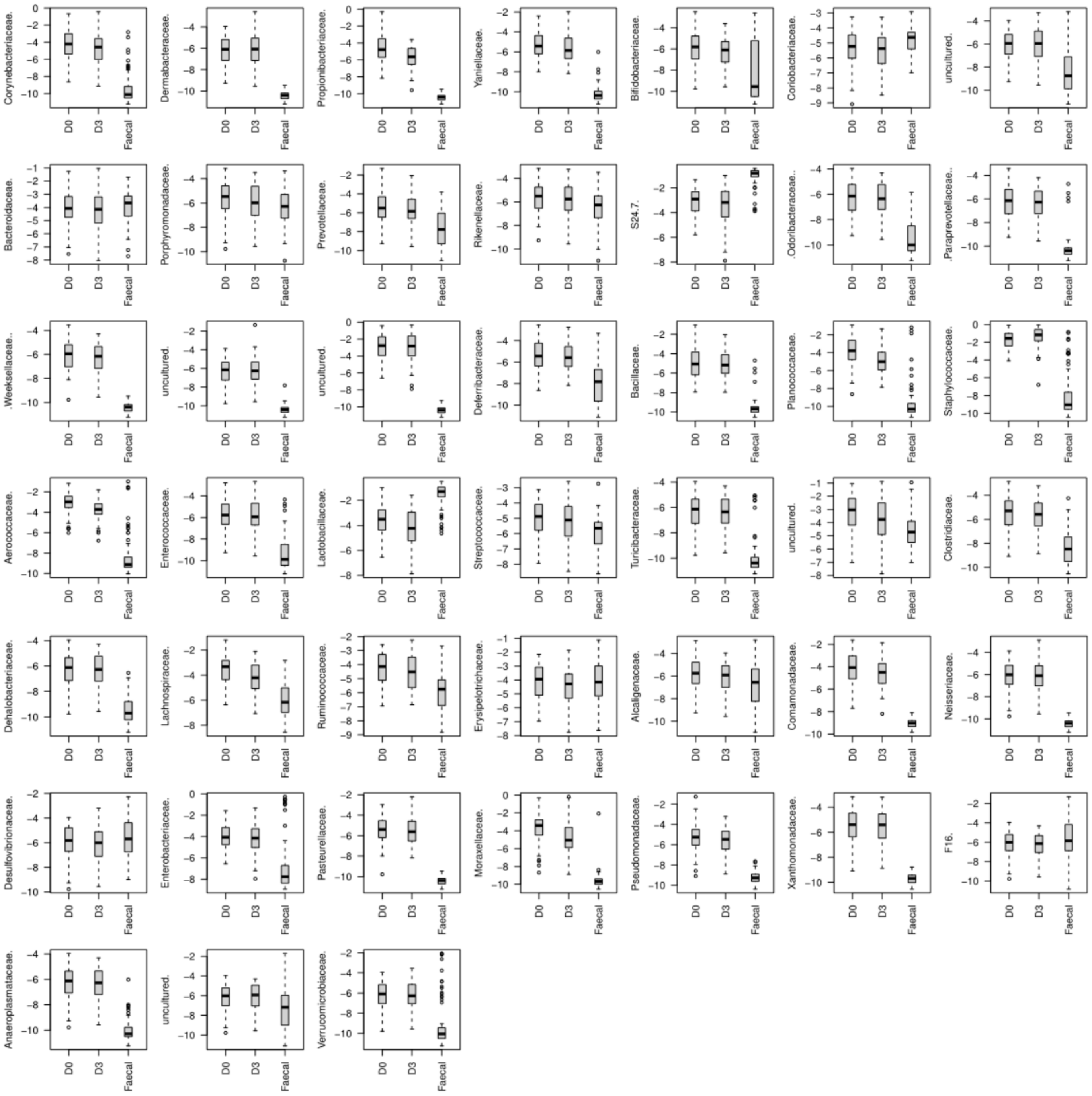


**b.**

Table 1: Linear Model of Staphylococcus and Aerococcus Diversity Associations

| Bacteria | Estimate | P.Value |
|---|---|---|
| Staphylococcus | -0.0751578 | 0.0682618 |
| Aerococcus | -0.1168848 | 0.0657272 |
